# The I-BAR domain protein baiap2l1a is required for protrusion and lateral elongation of epithelial microridge structures

**DOI:** 10.1101/2025.09.09.675250

**Authors:** Yasuko Inaba, Kimika Iwasaki, Aoi Nakamura, Shiro Suetsugu, Yasumasa Bessho

## Abstract

Microridges are laterally elongated membrane protrusions from the apical surface of epithelial cells. Microridges are arranged in striking maze-like patterns. They are found on various mucosal epithelia in many animals, including the skin of zebrafish, where they are required to maintain mucus on the skin surface. Recent studies have revealed molecular mechanisms of how microridiges formation involving actin and actin-regulatory proteins. However, the molecular mechanism that deforms epithelial membranes to create microridge protrusions remain unknown. We have found that one of the I-BAR domain proteins, baiap2l1a, which is known to regulate membrane curvature, is required for microridge morphogenesis. CRISPR/Cas9 knockdown showed that baiap2l1a mutant zebrafish had defects in microridge morphogenesis. Baiap2l1a mutant zebrafish had shorter and wider microridges than WT microridges. Baiap2l1a localized to microridges, and its localization proceeded microridge actin formation. Furthermore, the baiap2l1a I-BAR domain, which binds and curves membranes, was sufficient to localize to microridges in zebrafish skin cells. Structure/function experiments revealed that the I-BAR domain alone could partially rescued microridge length in baiap2l1a mutants. A 39 amino acid deletion in the I-BAR domain, which caused the loss of one α-helix according to AlphaFold2 simulations, is sufficient to impair microridge localization and failed to rescue microridge elongation in baiap2l1a mutants. These results suggest that the I-BAR domain is required for baiap2l1a microridge localization and function. Eps8like1a, a member of the Eps8 family proteins known as an actin capping and bundling protein genetically interacted with baiap2l1a in microridge elongation. Together, we found that the membrane curvature protein baiap2l1a plays an important role in generating microridges in zebrafish epithelia.

## Introduction

Microridges are laterally elongated protrusions from the apical surface of epithelial cells. Microridges are arranged in striking maze-like patterns. They are found on a variety of mucosal epithelia in many animals such as the human cornea and esophagus, and the skin of zebrafish epidermal cells, where they are required to maintain mucus on the skin surface ^12^. Early studies of microridges focused on describing microridge structure and distribution by electron microscope observation ^13^. Recent studies have addressed the molecular mechanism of microridge formation using zebrafish as a model ^4,5^.

Microridges contain both actin and keratin cytoskeletal filaments. Apicobasal polarity is important for microridge protrusion from the apical surface ^6^. Microridge formation is a dynamical process and composed of at least three steps. First, finger-like, actin-based protrusions called pegs protrude from the skin surface. Next, the pegs coalescence to form short microridges, a process dependent on the F-actin nucleator Arp2/3 and myosin-based contraction ^7,8^. Keratin intermediate filaments (IFs) contribute to microridge elongation, along with two cytolinker proteins periplakin and envoplakin ^9^. Once IFs invade microridges, microridges stabilize and elongate, using plakin-keratin interactions. Finally, microridges are then remodeled dynamically by fission and fusion to promote the formation of orderly microridge patterns ^8^.These recent studies have to reveal how cytoskeletal proteins promote microridge formation. However, the mechanisms that deform the apical membrane to form protrusive microridge structures remain unclear.

Bin-amphiphysin-Rvs (BAR) domain superfamily proteins are known as proteins that bind to the membranes and induce membrane curvature. The BAR domain superfamily is comprised of the classical BARs, F-BARs (Fes/CIP4 homology-BAR) and I-BARs (Inverse-BAR). The crystal structure of the BAR domains revealed a ‘banana’ shaped α-helical dimer that can bind to negatively charged membranes, causing the membrane curvature ^10,11^. BAR and F-BAR domain proteins drive positive membrane curvature and are involved in plasma membrane invaginations such as endocytosis, whereas I-BAR domain proteins bind the membrane and generate negative membrane curvatures to induce plasma membrane protrusions, such as filopodia and lamellipodia. There are five I-BAR domain proteins in mammals: IRSp53 (the insulin receptors substrate of 53 kDa), MIM (missing-in-metastasis suppressor 1), IRTKS (insulin receptor tyrosine kinase substrate; also known as BAIAP2L1), ABBA (actin-bundling protein with BAIAP2A homology) and BAIAP2l2 ^12,13,14^. I-BAR proteins participate in various morphogenetic events, such as microvilli formation, myoblast fusion, and dendritic spine formation ^15,16,17,18,19,20,21^.

In this study, we investigated how the membrane is deformed on the apical surface of zebrafish periderm cells to form microridges. Using a combination of live imaging, mutant analysis, simulations and structure-function studies, we found that one of the I-BAR proteins, baiap2l1a, has an important function in deforming the membrane to protrude the microridge structures. The actin-bundling protein Eps8like1a (EGFR pathway substrate 8 like1a) contributes to microridge morphogenesis and genetically interacts with baiap2l1a.

## Results

### 1. Baiap2l1a localizes in microridges and precedes actin filaments in microridge elongation

Mammals have five types of I-BAR domain proteins in mammals. A search of the Ensemble database revealed eight I-BAR domains proteins in zebrafish. Among these, an RNA-seq database shows that baiap2l1a and baiap2a were highly expressed in periderm cells ^22^. To determine if baiap2l1a localizes to microridges, we tagged baiap2l1a with GFP. Baiap2l1a localized to both pegs and microridges (Figure 1A-B). Interestingly, timelapse imaging of microridge morphogenesis revealed that baiap2l1a initially formed longer structures in a microridge pattern than actin, but actin later followed baiap2l1a localization, suggesting that baiap2l1a precedes actin during microridge morphogenesis (Figure 1C). These observations confirm that I-BAR domain protein baiap2l1a is a component of pegs and microridges, suggesting that it may play a role in microridge morphogenesis.

**Figure 1.**
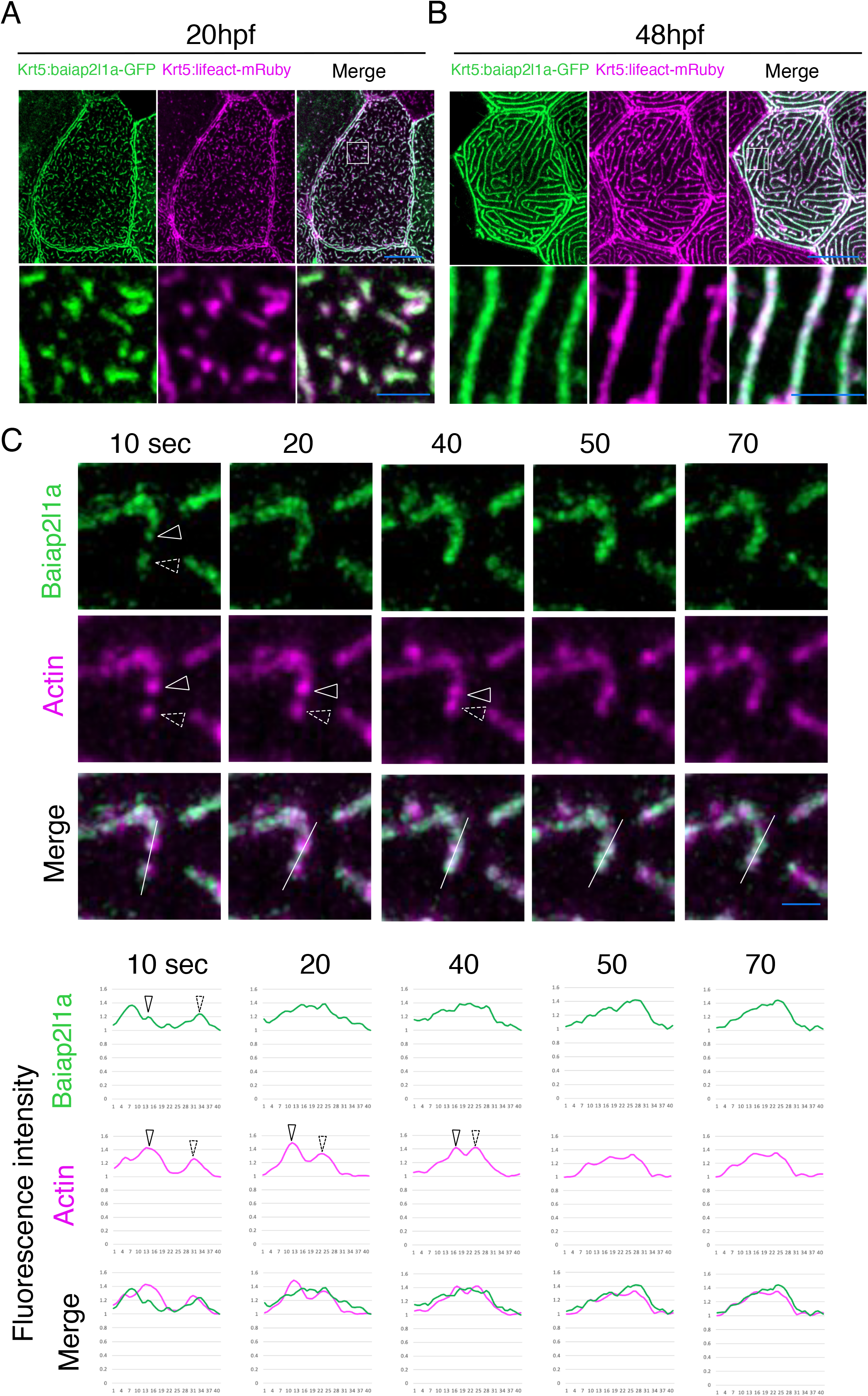
I-BAR domain protein baiap2l1a localized to pegs and microridges. (A) Confocal Airyscan images of periderm cells expressing baiap2l1a-GFP and Lifeact-mRuby in 20hpf zebrafish embryos. Boxes indicate zoomed-in regions of pegs and short microridges. (B) Confocal Airyscan images of periderm cells expressing baiap2l1a-GFP and Lifeact-mRuby in 48hpf zebrafish embryos. Boxes indicate zoomed-in regions of microridges. (C) Top: Sequential projections from a confocal Airyscan time-lapse movie of baiap2l1a-GFP and Lifeact-mRuby-expressing periderm cells during peg fused to short microridges (24hpf). Baiap2l1a structures appear to precede F-actin in developing protrusions. Bottom: Intensity plot of the line drawn in the top image showing baiap2l1a and actin signals. Arrowheads indicate the positions of fluorescence intensity shown in the top image. Scale bars: 10 µm (A-B) and 2 µm (C) zoomed images in A and B.

### 2. Baiap2l1a is required for protruding and elongation of microridges

To investigate if I-BAR domain proteins function in microridge morphogenesis, we knocked them down with CRISPR/Cas9. In F0 CRISPR-injected animals, we observed that baiap2l1a crispant embryos, but not baiap2a crispants, had defects in microridge formation (Supplement Fig.1), suggesting that baiap2l1a may contribute to microridge morphogenesis. To further examine the role of baiap2l1a in microridge elongation, we generated stable baiap2l1a zebrafish mutant lines by deleting several exons using the CRISPR/Cas9 system (Supplement Fig.1-2). We obtained two distinct baiap2l1a mutant alleles: one is an in-frame line 39 a.a. deletion in the I-BAR domain, as confirmed by sequencing data, and the other is a frameshift mutation (Figure.2A). To examine microridge morphology in heterozygous and homozygous mutants, we observed Krt5:lifeact-mRuby which label actin in periderm cells. Both mutant genotypes showed defects, with shorter microridges observed in both heterozygous and homozygous mutants, compared to WT (Figure 2B). Homozygous mutants had shorter microridges than heterozygotes (Figure 2C). To investigate the structures more closely, we performed Transmission Electron Microscopy (TEM) to measure the height and width of microridges. We found that microridges are shorter and wider in baiap2l1a homozygous mutants than WT. However, the distance between microridges was not significantly different from WT (Figure 2D-F). These results suggest that baiap2l1a contributes not only to lateral elongation but also to membrane curvature to promote protrusion from the cell surface during microridge morphogenesis. The in-frame allele (Δ39a.a. deletion in the I-BAR domain) also displayed defects in microridge length (Figure 2G-H). These results indicate that baiap2l1a is required for microridge protrusion and lateral elongation of microridges in periderm cells.

**Figure 2.**
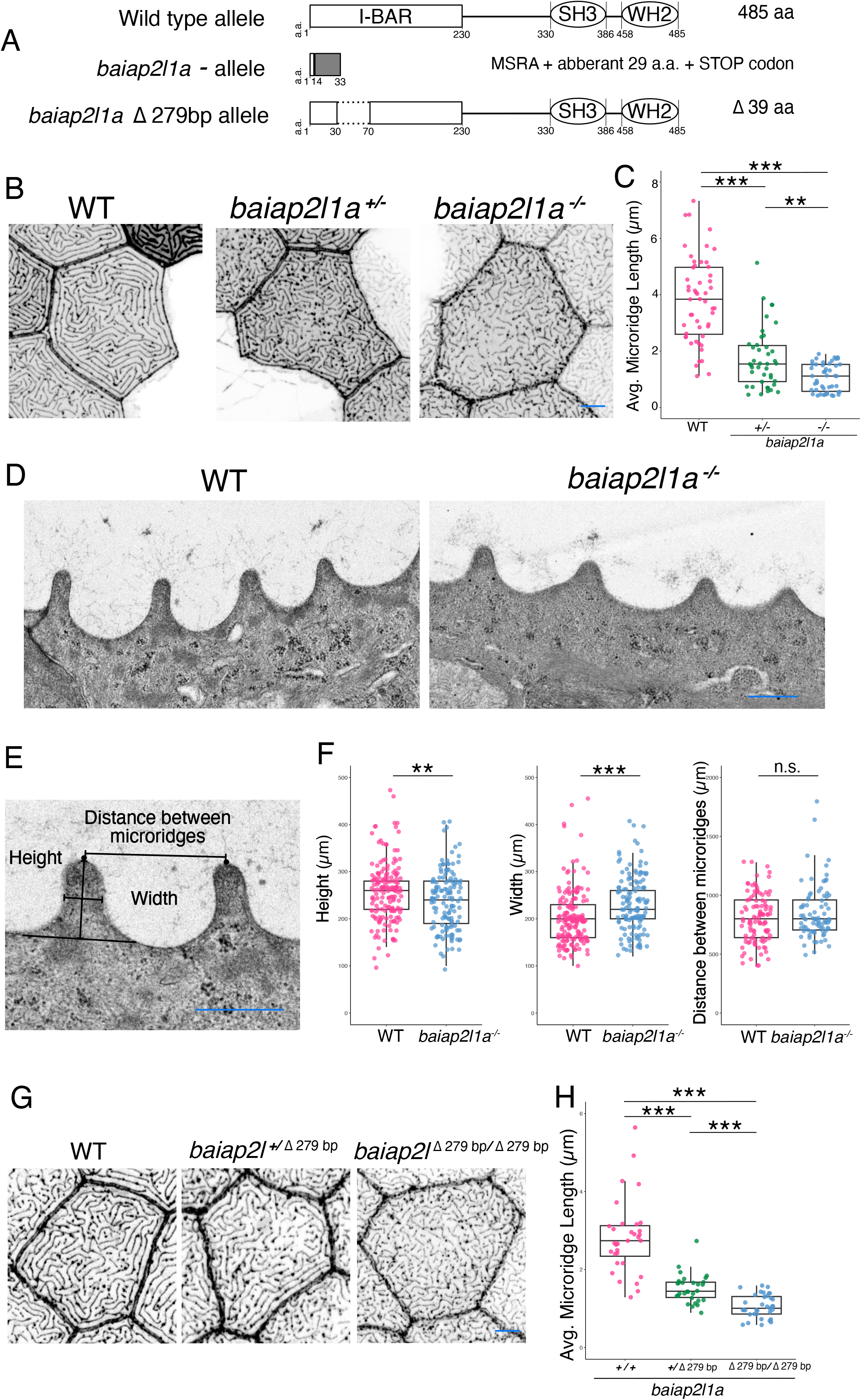
I-BAR domain protein baiap2l1a is required for peg protrusion and microridge elongation. (A) Baiap2l1a WT allele and mutant alleles. Expected protein structure from sequence. (B) Periderm cells expressing Lifeact-mRuby in WT and baiap2l1a mutants at 48hpf. Images were inverted. The high-intensity fluorescence appears black and low-intensity fluorescence is white. (C) Dot and box-and-whisker plot showing the average microridge length per cell at 48hpf in animals of the indicated genotypes. ^*^p<0.05, ^**^p<0.01, ^***^p<0.0001, the Wilcoxon rank-sum test. (D) TEM images of periderm cells in WT and baiap2l1a mutants at 48hpf. (E) The schematic indicating microridge height, width and distance between microridges used for quantification. (F) Dot and box-and-whisker plot showing the average microridge height, width, and distance between microridges per micoridge at 48hpf in animals of the indicated genotypes. ^*^p<0.05, ^**^p<0.01, ^***^p<0.0001, the t-est. (G) Periderm cells expressing Lifeact-mRuby in WT and baiap2l1a mutants at 48hpf. Images were inverted so that high-intensity fluorescence appears black and low-intensity fluorescence is white. (H) Dot and box-and-whisker plot showing the average microridge length per cell at 48 hpf in animals of the indicated genotypes. ^*^p<0.05, ^**^p<0.01, ^***^p<0.0001(Wilcoxon rank-sum test). Scale bars: 5 µm (B and G), 1 µm (D-E)

### 3. The I-BAR domain of baiap2l1a is sufficient for localization to microridges

I-BAR domain family proteins are composed of several domains: an N-terminal BAR domain that interacts with acidic phospholipids to curve the membrane, a central SH3 domain that binds to the actin regulatory complex, and an actin-binding Wiskott-Aldrich syndrome protein homology2 (WH2) motif at the C terminus ^12^. To investigate which baiap2l1a domain is required for baiap2l1a microridge localization in zebrafish periderm cells, we GFP-tagged each domain of baiap2l1a and expressed them in periderm cells in WT animals. The full-length baiap2l1a and I-BAR-SH3 domain (i.e.ΔWH2 domain) localized in microridges, but the SH-3-WH2 domain (i.e.ΔI-BAR domain) failed to localize in microridges. The WH2 domain, which directly binds to actin, also did not localize in microridges. Both the SH-3-WH2 and WH2 domains were localized throughout the cytoplasm. Interestingly, the I-BAR domain alone, which binds to the membrane, localized to microridges (Figure 3). These findings demonstrate that the baiap2l1a membrane binding I-BAR domain is required and sufficient for baiap2l1a microridge localization in zebrafish periderm cells.

**Figure 3.**
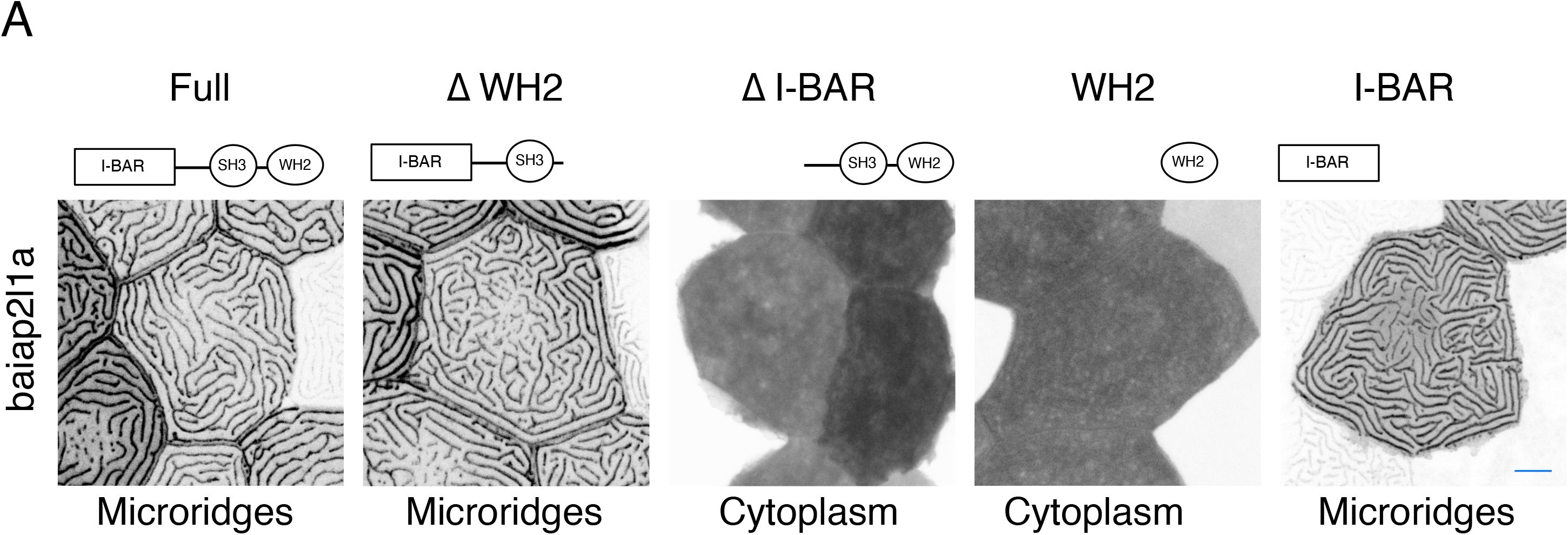
Baiap2l1a domain localization. (A) Periderm cells expressing baiap2l1a-GFP variants at 48hpf. Schematics indicate the domains in each variant. Images were inverted so that high-intensity fluorescence appears black and low-intensity fluorescence is white. Scale bars: 5 µm

### 4. The Baiap2l1a I-BAR domain is necessary and sufficient for microridge elongation

To investigate which baiap2l1a domain is required for microridge formation, we performed rescue experiments using several combinations of baiap2l1a domain constructs. First, we examined if the full-length baiap2l1a could rescue the baiap2l1a homozygote phenotype. We found that full-length baiap2l1a-injected cells could form elongated microridges and rescued the homozygous mutant microridge phenotypes (Figure 4A-B). Similarly, rescuing with the I-BAR-SH3 domain (i.e., ΔWH2 domain) could rescue the baiap2l1a homozygous mutant phenotype. However, the SH-3-WH2 domain (i.e., ΔI-BAR domain) could not rescue microridge elongation. Interestingly, the I-BAR domain alone could rescue microridge elongation partially, while the WH2 domain alone could not (Figure 4A-B). These results suggest that the I-BAR domain is required for microridge elongation, but the WH2 actin binding domain is not.

**Figure 4.**
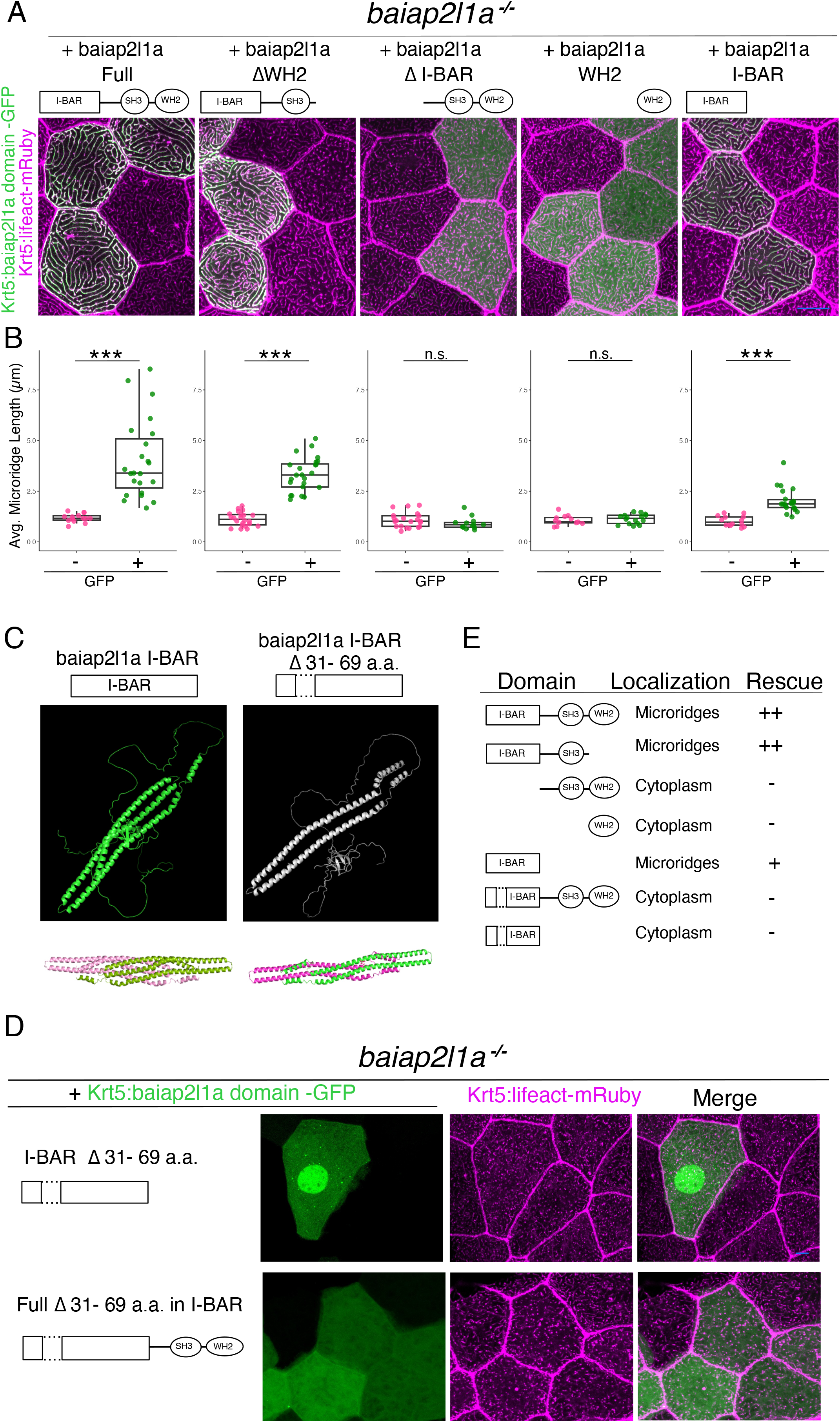
The Baiap2l1a I-BAR domain is partially rescues microridge elongation. (A) Lifeact-mRuby-expressing cells mosaically expressing baiap2l1-GFP variants in periderm cells at 48hpf, as indicated. Neighboring GFP-negative cells serve as internal controls and which are baiap2l1a mutant cells. (B) Dot and box-and-whisker plot showing the average microridge length per cell at 48hpf in GFP positive and negative cells. ^*^p<0.05, ^**^p<0.01, ^***^p<0.0001 (t-est). (C) Structure simulation of I-BAR domain using AlphoFold2. The structure below shows the dimerization of each I-BAR domain. (D) Lifeact-mRuby expressing cells mosaically expressing Baiap2l1a-I-BAR domain deletion GFP variants in periderm cells at 24hpf, as indicated. Neighboring GFP-negative cells serve as controls. (E) Summary of domain localization and the results from the rescue experiments. Scale bars: 10 µm (A) and 5 µm (D).

Since the I-BAR domain monomer consists of three α-helices that dimerize into an antiparallel structure to bind and curve the membrane, we expected that the I-BAR domain structure is important for their function in microridge morphogenesis^23 24,25^. To investigate the baiap2l1a I-BAR domain structure, we performed an AlphaFold2 simulation and found that the baiap2l1a I-BAR domain had three α-helices, as expected. However, the 39a.a. deletion in the baiap2l1a I-BAR domain led to the loss of one α-helix, resulting in a two-α-helix structure (Figure 4C). To investigate whether if motif can form dimers or not, we investigated the dimerization using an AlphaFold2 simulation. We found that the three-α-helix I-BAR domain formed straight dimers, while the two-α-helix could still form dimers but with a malformed shape (Figure 4C).

To investigate if the I-BAR intact structure is required for microridge morphogenesis, we performed rescue experiments. The intact I-BAR domain localize to microridges and rescued microridge length in baiap2l1a homozygous mutants, but the two-α-helix I-BAR domain lost its microridge localization, only localizing to the cytoplasm, and failed to rescue the microridge elongation in baiap2l1a homozygous mutants (Figure 4D). We next investigated if the other domains could compensate for the two-α-helix I-BAR domain by making a full-length baiap2l1a with the 39a.a. deletion in the I-BAR domain. This construct did not localize to microridges and failed to rescue the microridge elongation in homozygous mutants (Figure 4D). These results suggest that the other baiap2l1a domains cannot compensate for the two-α-helix I-BAR domain, and the intact three-α-helix structure is necessary for baiap2l1a localization and function in microridges.

### 5. Baiap2l1a and Eps8l1a exhibit a genetical interaction in micorirdge elongation

To investigate what other proteins work with baiap2l1a in microridge elongation, we focused on eps8l1a. Eps8 is known as an actin-bundling and capping protein. It has been shown that baiap2l1a, which is IRTKS in humans, binds to eps8 to elongate microvilli in the intestinal epithelium. Furthermore, Eps8l1a is highly expressed in zebrafish periderm cells, which have microridges.

To examine the role of eps8l1a, we first observed its localization and found that it localized in microridges. Eps8l1a localization was independent of baiap2l1a (Supplement Fig.4). Baiap2l1a microridge localization was also independent of eps8l1a (Supplement Fig.4). Next, we created eps8l1a mutant zebrafish. Eps8l1a mutants did not have microridge defects (Supplement Fig.5). Finally, we next tested if eps8l1a and baiap2l1a have interact genetically in microridge morphogenesis. Indeed, animals that were heterozygous for baiap2l1a and homozygote for eps8l1a formed shorter microridges than baiap2l1a heterozygotes. Furthermore, eps8l1a and baiap2l1a double mutants formed shorter microridges than baiap2l1a mutants alone (Fig.5). These results suggests that eps8l1a and baiap2l1a genetically interact in microridge morphogenesis.

## Discussion

This study has uncovered a mechanism by which I-BAR domain proteins contribute to the formation of protrusive, laterally elongated microridges on mucosal epithelial membranes. Recent investigations have addressed the molecular mechanisms underlying microridge formation, including the involvement of actin and other cytoskeletal elements. However, the molecular mechanism by which proteins deform membranes were unknown.

We found that an I-BAR domain proteins, baiap2l1a, a homologue of IRTKS in humans, plays an important role in microridge morphogenesis. BAR domain proteins are membrane curvature proteins. Specifically, I-BAR domains binds the membrane to induce negative membrane curvatures to form protrusive structures, such as filopodia and lamellipodia. IRTKS promotes the growth of epithelial microvilli, along with eps8, in intestinal epithelial cells ^26^.

We found that baiap2l1a localizes to microridges, and time-lapse imaging revealed that baiap2l1a precedes actin in microridges in zebrafish epithelial cells (Figure 1A-C). In baiap2l1a mutants, microridge formation is disrupted, resulting in wider and shorter structures compared to WT microridges, as observed by TEM at 48hpf (Figure 2D-F). Microridge lateral length was also shorter in mutants (Figure 2B-C).

Microridges form periodic maze-like patterns in mucosal epithelia cells. A recent study showed that local non-muscle myosin Ⅱ(NMⅡ) activity in the apical cortex promote microridge fission and fusion, contributing to the formation of a regularly spaced microridge pattern ^7^. However, mechanisms that regulate the width of microridges themselves are unknown. Baiap2l1a mutant embryos had wider microridges, suggesting that the regulation of membrane deformation is important for forming the regular width of microridges. Further investigations are required to reveal further mechanisms of microridge width regulation.

Baiap2l1a consists of three domains: an N-terminal I-BAR domain that binds and curves the membrane, a central SRC homology domain 3 (SH3), which in other proteins can bind to actin-interacting proteins, and a WH2 domain at the C-terminus, which directly binds to actin. By analyzing the role of each domain in baiap2l1a localization, we found that the I-BAR domain alone is sufficient for baiap2l1a to localize to microridges, similar to the baiap2l1a full-length protein. Deletion of the WH2 domain showed no effect on baiap2l1a microridge localization. However, deletion of the I-BAR domain impaired baiap2l1a localization in microridges (Figure 3). Since the I-BAR domain is known to bind to the membrane, these results suggest that actin interactions are not necessary for baiap2l1a microridge localization. Instead, baiap2l1a interacts with the membrane through its I-BAR domain. Similarly, the human homologue IRTKS localizes to the tips of microvilli in the intestinal epithelium, and the IRTKS I-BAR domain alone exhibits similar localization in culture cells ^26^.

To investigate if interaction with the membrane is sufficient for baiap2l1a’s function in microridge morphogenesis, we performed rescue experiments with several domain deletion constructs. In baiap2l1a mutants, the full-length baiap2l1a rescued the microridge length. Similarly, the baiap2l1a construct without the WH2 domain (I-BAR and SH3 domains are still intact) rescued microridge length. However, the construct without the I-BAR domain did not rescue microridge length. Interestingly, the I-BAR domain alone partially rescued the microridge length (Figure 4A). These results suggest that generating membrane curvature through the I-BAR domain is important for baiap2l1a-induced microridge elongation. Since the I-BAR domain alone rescued the microridge length partially, we expected that other proteins work together with baiap2l1a in microridge elongation.

Eps8 is as an actin-capping and bundling protein. It was previously reported that IRTKS and eps8 work together to form and elongate microvilli in the intestinal epithelium. Therefore, we investigated if this protein interaction also contributes to the formation of microridges. We found that eps8l1a single mutants did not have microridge length defects at 48hpf. However, baiap2l1a heterozygous; eps8l1a homozygous mutants had shorter microridges than microridges in heterozygous baiap2l1a mutant alone. Furthermore, eps8l1a; baiap2l1a double mutants had shorter microridges compared to baiap2l1a single mutant microridges (Figure 5). These results indicate that baiap2l1a and has a genetic interaction with esp8l1a in microridge morphogenesis and may function upstream of eps8l1a.

**Figure 5.**
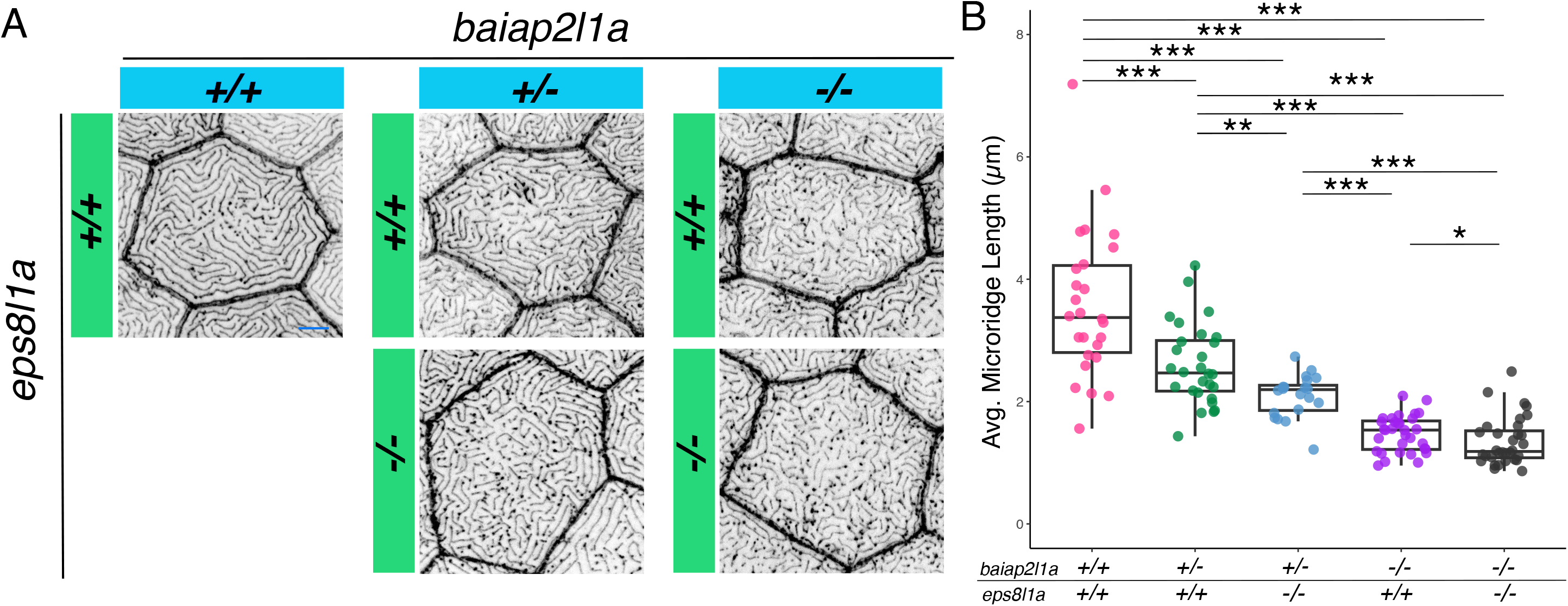
Baiap2l1a and eps8l1a have a genetic interaction in microridge morphogenesis. (A) Periderm cells expressing Lifeact-mRuby with the indicated genotypes at 48hpf. Images were inverted so that high-intensity fluorescence appears black and low-intensity fluorescence is white. (B) Dot and box-and-whisker plot showing the average microridge length per cell at 48hpf in animals of the indicated genotypes. ^*^p<0.05, ^**^p<0.01, ^***^p<0.0001 (Wilcoxon rank-sum test). Scale bars: 10 µm.

The structure of I-BAR domain proteins is important for its binding activity to the membrane. We obtained an in-frame mutant fish line with CRISPR/Cas9 mutagenesis, which is expected to have 39a.a. deletion. Interestingly, this deletion impairs microridge morphogenesis, since homozygous mutants with 39a.a. deletion had showed shorter microridges than WT, similar to deletions of the entire protein. From structural analysis using AlphoFold2 simulation, we found that the 39a.a. deletion results in the loss of one α-helix in the I-BAR domain. In WT, baiap2l1a, the I-BAR domain has three α-helixes, whereas only two α-helix structures from in the 39a.a. deletion, which impaired microridge elongation. These results suggest that the intact I-BAR domain structure is important for binding to the membrane to elongate microridge structures. Collectively, these results suggest that an intact I-BAR domain is required for baiap2l1a microridge localization and function in microridges.

Interestingly, baiap2l1a localized not only in microridges, but also in peg structures, suggesting its potential role in peg membrane protrusion, in addition to the elongation of protruding microridge membranes. Further investigation may reveal the detail mechanism by which I-BAR domain proteins contribute to membrane protrusion in mucosal epithelia.

---

**Supplementary Figure 1.**
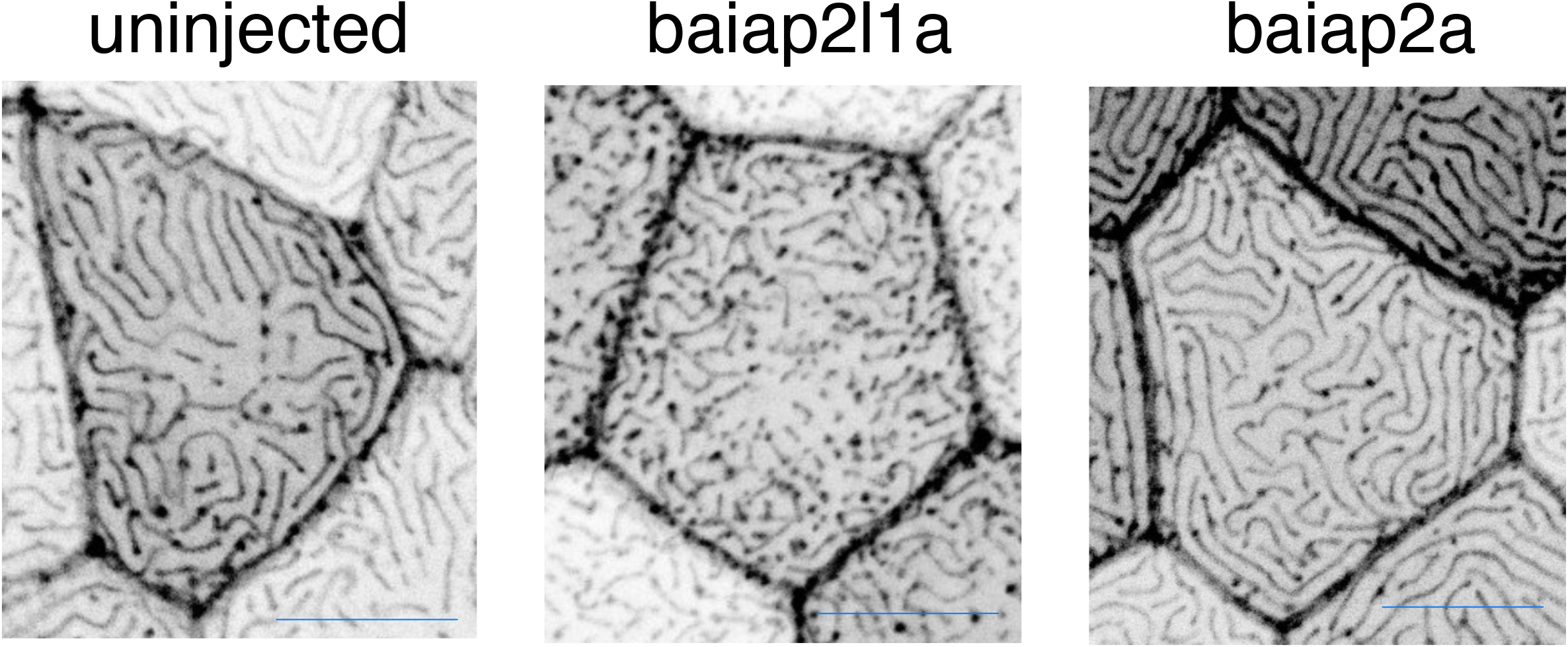
Baiap2l1a and baiap2a gRNA injected F0 embryos. Uninjected embryos and embryos injected with *baiap2l1a* and *baiap2a* gRNAs at 48 hpf. Scale bars: 10 µm (A) and 5 µm (D)

**Supplementary Figure 2.**
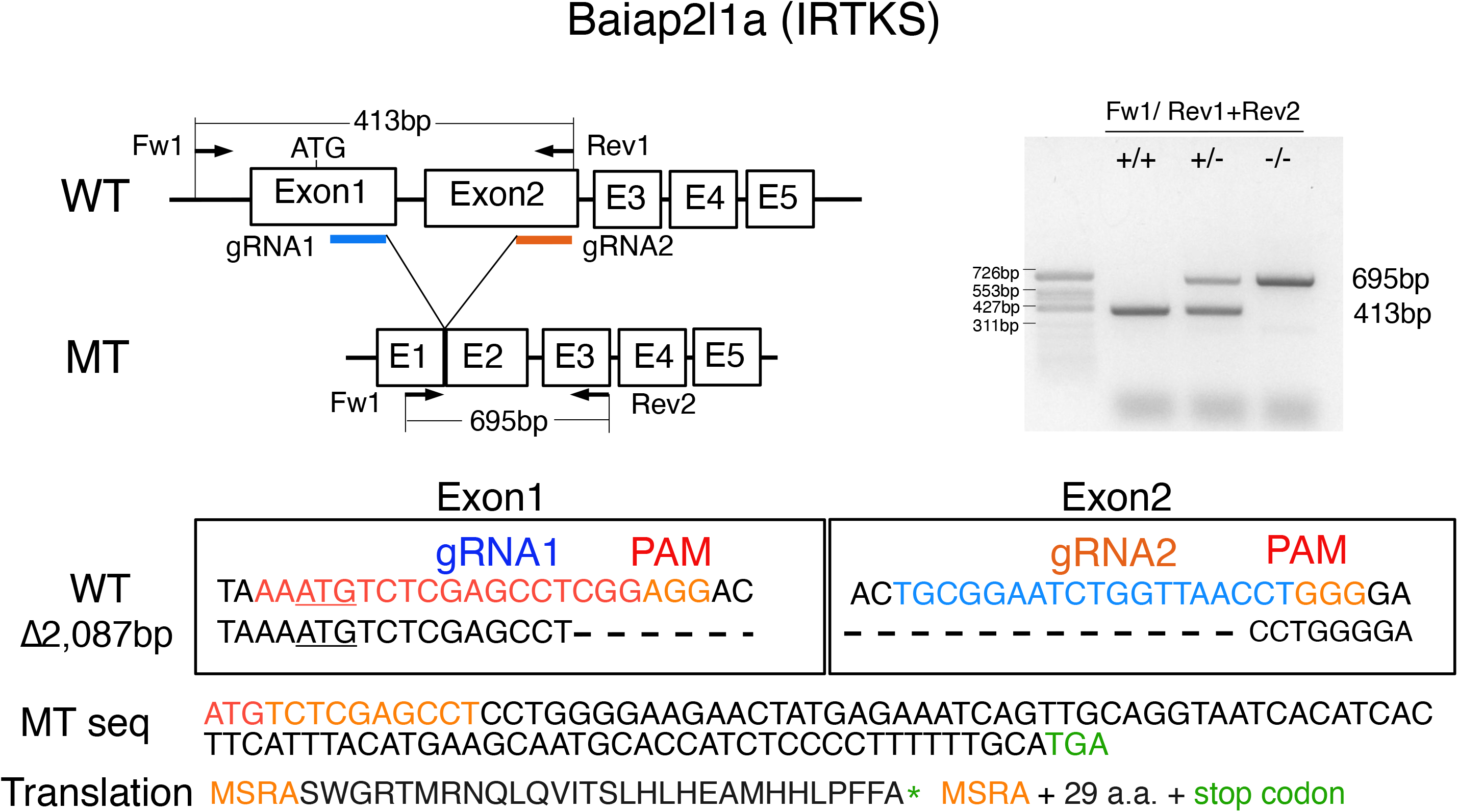
Baiap2l1a deletion mutant. The *baiap2l1a* gene targeting strategy. Sequence shows the deletion site. Right panel shows the PCR analysis of genomic DNA isolated from adult fish fins with the indicated genotypes. Primers Fw1 and Rev1 were used to detect WT alleles (413bp) and primers Fw1 and Rev2 were used to detect mutant alleles (695bp).

**Supplementary Figure 3.**
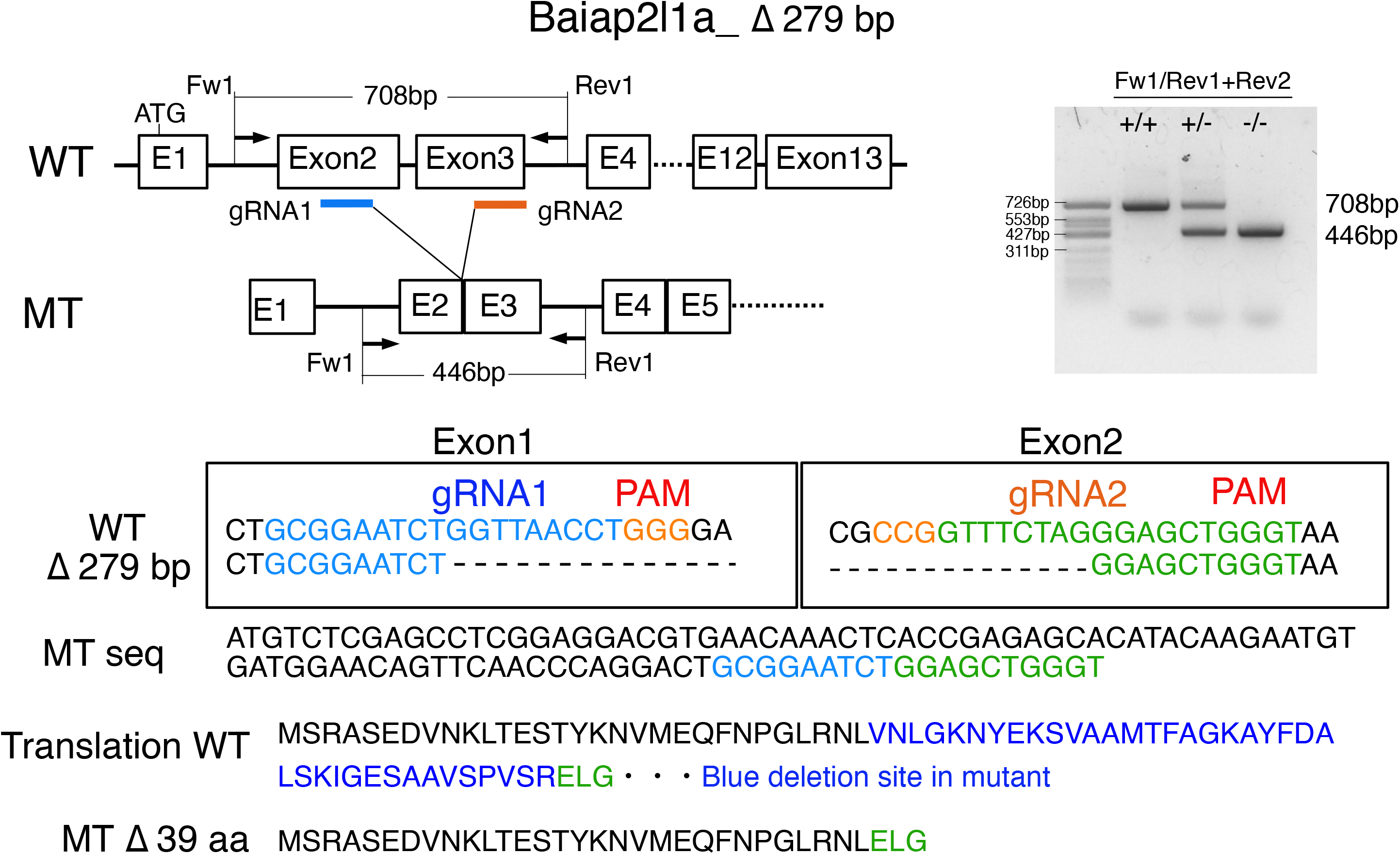
Δ39a.a. baiap2l1a deletion mutant. The *baiap2l1a* gene targeting strategy. Right panel shows the PCR analysis of genomic DNA isolated from adult fish fins with the indicated genotypes. Primers Fw1 and Rev1 were used to detect WT alleles (708 bp) and primers Fw1 and Rev2 were used to detect mutant alleles (446 bp). The sequence below shows the deletion site which is an in-frame deletion.

**Supplementary Figure 4.**
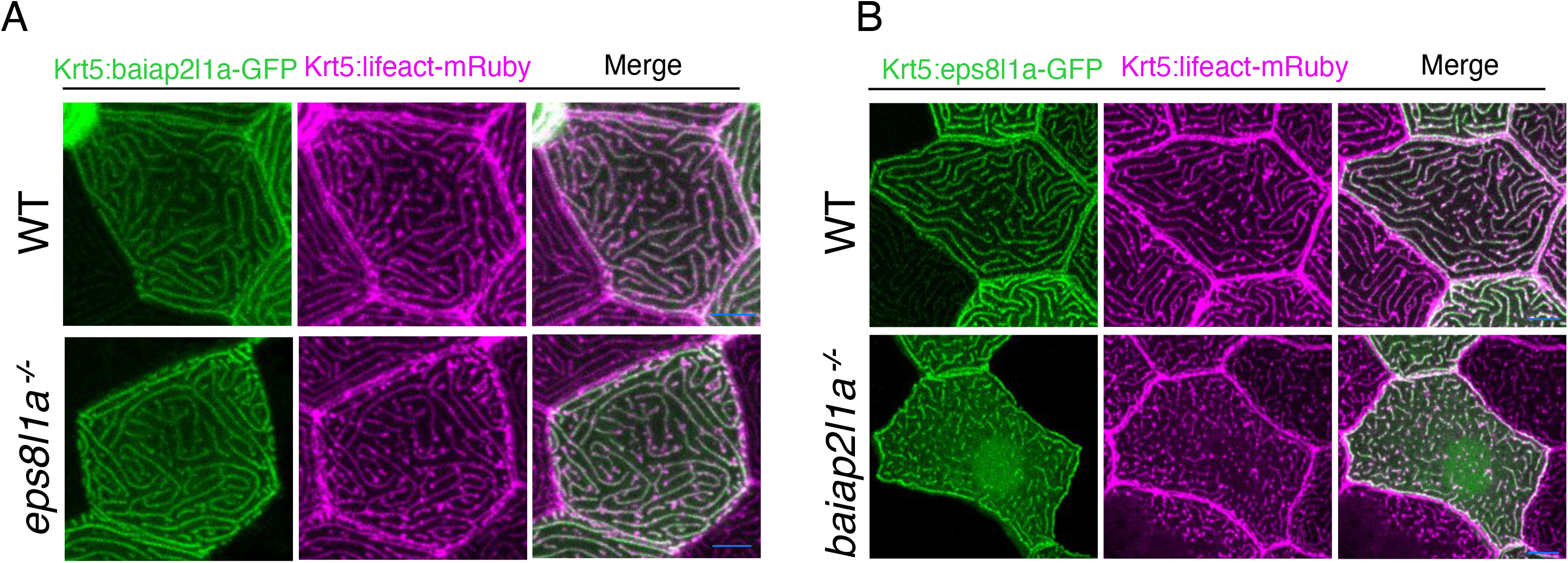
Baiap2l1a and eps8l1a localization is independent of each other. (A) Periderm cells expressing Baiap2l1a-GFP and Lifeact-mRuby in WT and eps8l1a^-/-^ periderm cells. Baiap2l1a localized to microridges in both WT and eps8l1a^-/-^ periderm cells at 48hpf. (B) Periderm cells expressing eps8l1a-GFP and Lifeact-mRuby expressing in WT and baiap2l1a ^-/-^ periderm cells. Eps8l1a localized to microridges in WT and pegs/short microridges, nucleus and cytoplasm in baiap2l1a^-/-^ periderm cells at 48hpf. Scale bars: 10 µm.

**Supplementary Figure 5.**
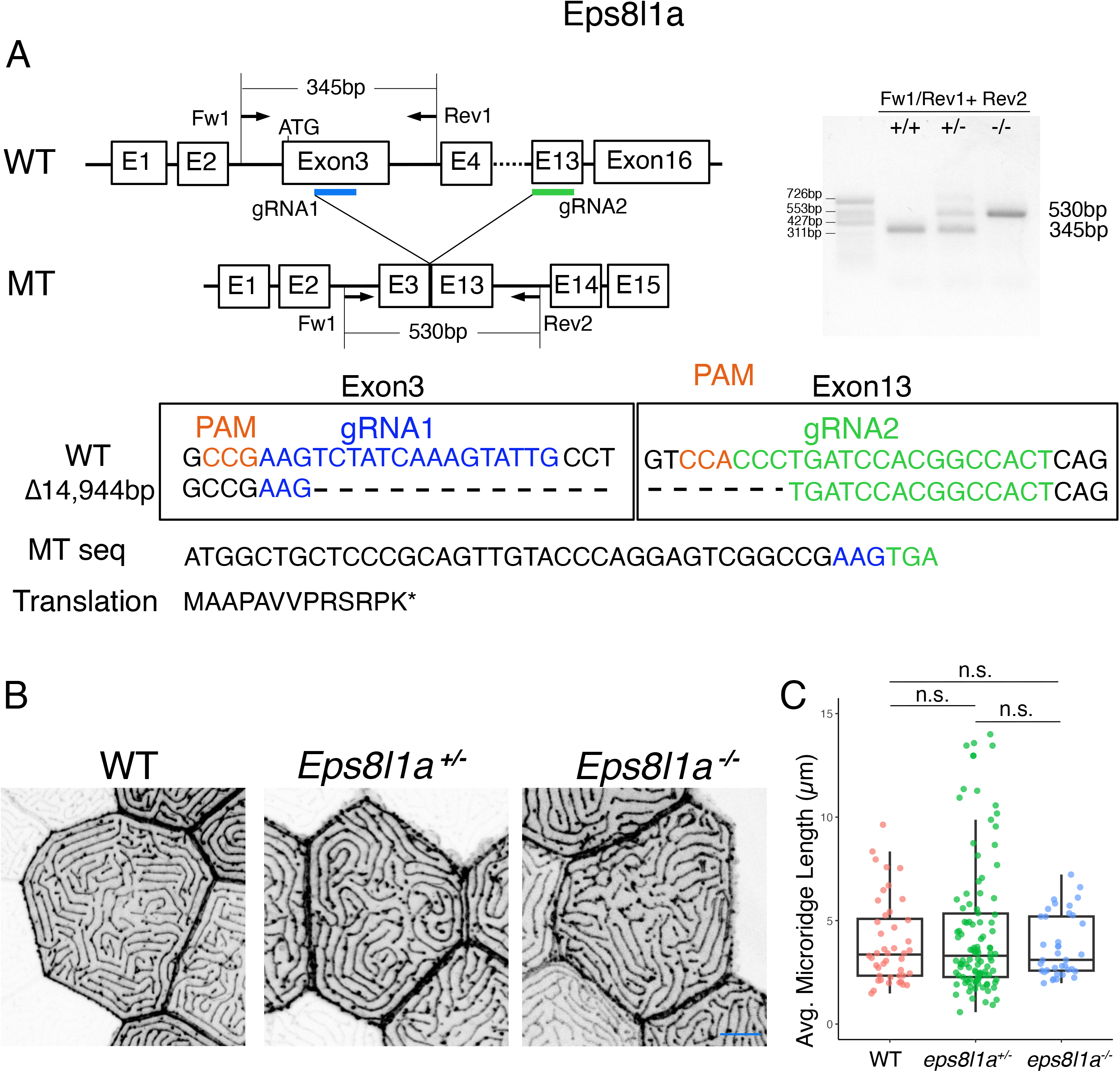
Eps8l1a single mutant does not affect microridge morphogenesis. (A) The *eps8l1a* gene targeting strategy. Right panel shows the PCR analysis of genomic DNA isolated from adult fish fins with the indicated genotypes. Primers Fw1 and Rev1 were used to detect WT alleles (345 bp) and primers Fw1 and Rev2 were used to detect mutant alleles (530 bp). The sequence below shows the deletion site. (B) Periderm cells expressing Lifeact-mRuby in WT and eps8l1a mutants at 48hpf. Images were inverted so that high-intensity fluorescence appears black and low-intensity fluorescence is white. (C) Dot and box-and-whisker plot showing the average microridge length per cell at 48hpf in animals of the indicated genotypes. ^*^p<0.05, ^**^p<0.01, ^***^p<0.0001 (Wilcoxon rank-sum test). Scale bars: 10 µm.

## Material and methods

### Zebrafish

Zebrafish (*Danio rerio*) were maintained at 28°C on a 14-hour light/10-hour dark cycle. Embryos were raised at 28.5°C in embryo water (0.3 g/L lnstant Ocean salt, 0.1% methylene blue). All experimental procedures were approved by the Chancellor’s Animal Research Care Committee at NAIST.

### CRISPR/Cas9 mutagenesis

To generate guide RNAs (gRNAs), we used the ‘short oligo method to generate gRNA’, as previously described (<u>Talbot and Amacher, 2014</u>). Two Cas9 binding sites were selected for each gene. *baiap2l1a* targeting sequences were located in exons 1 and 2 for mutant, and exons 2 and 3 for the 39 a.a. deletion line. *Eps8l1a* targeting sequences were in exons 3 and 13. The DNA template was PCR-amplified to make a product containing a T7 RNA polymerase promoter, the gene targeting sequence, and a gRNA scaffold sequence. PCR products were used as a template for RNA synthesis with T7 RNA polymerase (New England Biolabs) and purified (QIAGEN RNA purification kit) to generate gRNAs. Injection mixes contained Cas9 protein (1 mg/mL; FASMAC), gRNAs (0.5–1 ng/µL), and 300 mM KCI. Injection mixes were incubated on ice for 15 mins before injection. Embryos were injected at the 1-cell stage with 2–5 nL of injection mix. To identify germline founders, F0 fish was crossed with wild-type fish and the resulting 48hpf embryos were collected for PCR genotyping. Founder progenies were raised to adulthood to establish stable mutant lines.

### Plasmids

Plasmids were constructed using the HiFi DNA assembly (NEB). Primer sequences are listed in the Key Resources Table. Krt5:baiap2l1a-I-BARΔ39 a.a. in full-GFP and Krt5:baiap2l1a-I-BAR Δ39 a.a.-GFP transgenes were created using PCR from Krt5:baiap2l1a-GFP or Krt5:baiap2l1a-IBAR domain-GFP plasmids with SuperFi DNA Polymerase (Invitrogen). PCR products were gel extracted and transformed, and the selected colonies were sequenced. The following plasmids were previously described: Krt5-Lifeact-GFP and Krt5-Lifeact-mRuby (<u>van Loon et al., 2020</u>).

### Mounting embryos for live imaging

Live zebrafish embryos were anaesthetized with ∼0.2 mg/mL MS-222 (tricaine) in system water prior to mounting. Embryos were embedded in 1.2% LM agarose (nacalai tesque) on a glass bottom dish filled with 0.2 mg/mL MS-222 solution.

#### Microscopy

Confocal imaging was performed using an LSM 980 microscope with Airyscan (Carl Zeiss) using a 40x oil objective (NA = 1.3).

### Image analysis and statistics

Image analysis was performed with FIJI (<u>Schindelin et al., 2012</u>). For display purposes, confocal z-stack images were projected (maximum intensity projection) and brightness and contrast were optimized. The Image Stabilizer plugin was used to adjust for cell drift. An automated pipeline implemented in FIJI was used to analyze the average microridge length per cell, as previously described ^9^(<u>van Loon et al., 2020</u>).

Statistical analyses and graphs were generated with RStudio. Details of statistics for each experiment are listed in the figure legends.

### Simulation

The amino acid seq of Zebrafish WT I-BAR domain and Δ39a.a I-BAR domain used for the structure prediction thought Alphsfold2 as describe previously^27^.

### TEM

The samples were fixed with 2 % paraformaldehyde (PFA) and 2 % glutaraldehyde (GA) in 0.1 M cacodylate buffer pH 7.4 at 4 °C overnight. Samples were then postfixed with 2 % osmium tetroxide (OsO_4_) in 0.1 M cacodylate buffer. The samples were dehydrated in graded ethanol solutions (50 %, 70 %, 90 %, and anhydrous ethanol). Following dehydration, the samples were infiltrated with propylene oxide (PO) 2 (Quetol-812; Nisshin EM Co., Tokyo, Japan). The samples were then transferred to a fresh 100 % resin, and were polymerized at 60 °C for 48 h. The polymerized resins were ultra-thin sectioned at 70 nm with a diamond knife on an ultramicrotome (Ultracut UCT; Leica,Vienna, Austria), and the sections were mounted on copper grids. The grids were stained with 2 % uranyl acetate at room temperature, washed with distilled water, and then secondary-stained with Lead stain solution (Sigma-Aldrich Co., Tokyo, Japan). The grids were observed by a transmission electron microscope (JEM-1400Plus; JEOL Ltd., Tokyo, Japan) at an acceleration voltage of 100 kV. Digital images (3296 × 2472 pixels) were capture with a CCD camera (EM-14830RUBY2; JEOL Ltd.).

## Acknowledgements

We thank Alvaro Sagasti for careful reading of the manuscript and discussion. We also thank members of the Bessho lab for comments on the manuscript, Maiko Yokouchi and Yuka Ueda for excellent fish care, and NAIST Life Science Collaboration Center (Lisco) microscopy core.

